# Comparing low-pass sequencing and genotyping for trait mapping in pharmacogenetics

**DOI:** 10.1101/632141

**Authors:** Kaja Wasik, Tomaz Berisa, Joseph K. Pickrell, Jeremiah H. Li, Dana J. Fraser, Karen King, Charles Cox

## Abstract

Low pass sequencing has been proposed as a cost-effective alternative to genotyping arrays to identify genetic variants that influence multifactorial traits in humans. For common diseases this typically has required both large sample sizes and comprehensive variant discovery. Genotyping arrays are also routinely used to perform pharmacogenetic (PGx) experiments where sample sizes are likely to be significantly smaller, but clinically relevant effect sizes likely to be larger. To assess how low pass sequencing would compare to array based genotyping for PGx we compared a low-pass assay (in which 1× coverage or less of a target genome is sequenced) along with software for genotype imputation to standard approaches. We sequenced 79 individuals to 1× genome coverage and genotyped the same samples on the Affymetrix Axiom Biobank Precision Medicine Research Array (PMRA). We then down-sampled the sequencing data to 0.8×, 0.6×, and 0.4× coverage, and performed imputation. Both the genotype data and the sequencing data were further used to impute human leukocyte antigen (HLA) genotypes for all samples. We compared the sequencing data and the genotyping array data in terms of four metrics: overall concordance, concordance at single nucleotide polymorphisms in pharmacogenetics-related genes, concordance in imputed HLA genotypes, and imputation r^2^. Overall concordance between the two assays ranged from 98.2% (for 0.4× coverage sequencing) to 99.2% (for 1× coverage sequencing), with qualitatively similar numbers for the subsets of variants most important in pharmacogenetics. At common single nucleotide polymorphisms (SNPs), the mean imputation r^2^ from the genotyping array was 90%, which was comparable to the imputation r^2^ from 0.4× coverage sequencing, while the mean imputation r^2^ from 1× sequencing data was 96%. These results indicate that low-pass sequencing to a depth above 0.4× coverage attains higher power for trait mapping when compared to the PMRA.

## Introduction

Research in human genetics relies on efficiently profiling the genome of large numbers of individuals. A number of approaches can be used for this, usually trading off comprehensiveness (i.e. the fraction of the genome that is measured) with cost. By far the most commonly-used approach is the genotyping array, in which a set of known polymorphisms (usually around 500,000-2,000,000) is measured. This technology is inexpensive (currently on the order of tens to hundreds of dollars), but the set of genetic variants profiled is a small number of all known variants, and the technology does not allow for the detection of new (for example rare or population-specific) genetic variants. Genotyping arrays are commonly used for pharmacogenetics (PGx) studies where typically sample numbers are more limited, but inclusion of PGx focused variants on the arrays makes them suitable tools for screening the genome for markers associated with efficacy and adverse events.

The technological alternative to genotyping technology is sequencing technology, in which specific polymorphisms are not targeted for analysis, but rather the entire genome is sampled with some average depth of coverage. As sequencing costs have dropped, low-pass sequencing (for our purposes, which we will define as sequencing in which the average coverage of the genome is equal to or lower than 1×) becomes an appealing alternative to genotyping (CONVERGE consortium et al., 2015; Gilly et al., 2017; Pasaniuc et al., 2012). As an intuition for why this approach is useful, note that a human sample sequenced at 0.4× coverage is expected to have a single sequencing read covering each of around 30 million genetic variants identified in the 1000 Genomes Project (Auton et al., 2015), while a genotyping array obtains measurements (albeit somewhat less noisy measurements) at two orders of magnitude fewer sites.

In this paper, we directly compare genotyping results from low-pass sequencing to a commonly used genotyping array, the Affymetrix Axiom Biobank Precision Medicine Research Array (PMRA). Two types of metrics are relevant for this comparison. One is simply the genome-wide coverage of the assay, which we measure using average imputation quality. The other is genotyping quality at particular genetic variants of interest. We were particularly interested in applications to PGx—the identification of genetic variants that influence drug response. In this application, genetic variants in the MHC and genes involved in drug metabolism (so-called “ADME” genes, for absorption, distribution, metabolism, and excretion) are known to be particularly relevant. We thus considered these separately.

## Results

We selected 79 individuals to be both genotyped and sequenced. Each individual was genotyped on the Affymetrix Axiom Biobank PMRA, and sequenced by Gencove, Inc. to an average of 1× coverage using the Illumina HiSeq 4000 platform with paired-end 150 base pair reads. Sequencing reads were then sampled at random to obtain an average of 0.8×, 0.6×, and 0.4× coverage of the genome (Methods).

We then performed genotyped imputation of genetic variants in the 1000 Genomes Phase 3 release. This imputation for was performed using minimac2 (for the genotyping array data) or Gencove’s loimpute software v0.18 (for the low-pass sequencing data, see Methods for details). Both the unimputed PMRA data and the imputed low-pass sequencing data were then used to impute HLA genotypes using HIBAG (Zheng et al., 2014).

The relevant metrics to use when comparing the two technologies depend on the downstream use cases. Specifically, if an investigator is interested in identifying genetic variants associated with a trait but has no *a priori* knowledge of where in the genome such variants are likely to be located, then the relevant metric is the average correlation between imputed genotype calls and true genotypes. On the other hand, if the investigator knows that specific variants are most likely to be relevant to the trait of interest, then the relevant metric is the concordance between the technologies at those specific sites. Since in PGx applications there are some specific genes and variants of interest, we computed metrics in both of these classes.

### Overall genotype concordance

We first examined the overall concordance between the genotyping arrays and imputed sequences at different depths. To do this, we removed genotypes imputed with low confidence (with less than 90% posterior probability on a single genotype), and assessed the concordance between the two platforms, averaging across individuals, using metrics from the draft guidance of the United States Food and Drug Administration. These metrics measure concordance for variants present and absent in a reference genome—a “positive percent agreement” (PPA) for variants that are different from the reference and a “negative percent agreement” (NPA) for variants that match a reference genome. For our purposes we considered the genotypes from the PMRA as “truth”; in this case the PPA ranged from 98.2% for 0.4× coverage sequencing to 99.2% for 1× coverage sequencing, while the NPA ranged from 99.8% for 0.4× coverage to 99.9% for 1× coverage.

**Table 1:** Genotype concordance between genotyping and sequencing platforms. In all cases the genotyping array was treated as ‘Truth’.**Positive % Agreement (PPA)**– The percent of non-reference calls in the Truth dataset detected by Test, ignoring no calls in Test. (True Positives / True Positives + False Negatives).**Negative % Agreement (NPA)** – The percent of reference calls in the Truth dataset detected by Test, ignoring no calls in Test. (True Negatives / True Negatives + False Positives).No Calls– Count of **No Calls** in test that were variant in Truth. No calls are averaged across all 79 individuals.

| Comparison | PPA (%) | NPA (%) | No Calls (Average) |
| --- | --- | --- | --- |
| Accuracy, .4x vs PMRA | 98.22% | 99.82% | 2535 |
| Accuracy, .6x vs PMRA | 98.76% | 99.85% | 1848 |
| Accuracy, .8x vs PMRA | 99.01% | 99.86% | 1508 |
| Accuracy, 1x vs PMRA | 99.19% | 99.88% | 1251 |

### Genotype concordance at ADME genes

We then specifically compared the concordance between the genotypes at variants in ADME genes as defined by Hoverlson et al. (Hovelson et al., 2017). There were 216 such variants that were directly genotyped on the PMRA. We thus computed the same concordance metrics specifically at these 216 variants. For these analyses we excluded low-confidence genotype calls from the low-pass sequencing data; the percentage of excluded calls range from 1.6% of genotype calls in the 0.4× data down to 0.8% of genotype calls in the 1× data.

Concordance results are presented in Figure 1A. At common variants (where the minor allele is present in more than five copies in the sample, corresponding to a minor allele frequency over 3%), PPA ranged from 98.5% (for 0.4× coverage) up to 99.4% (for 1× coverage). The lowest concordance metric was the PPA at rarer variants, which ranged from 82.1% (for 0.4× coverage) to 95.2% (for 1× coverage).

**Figure 1:**
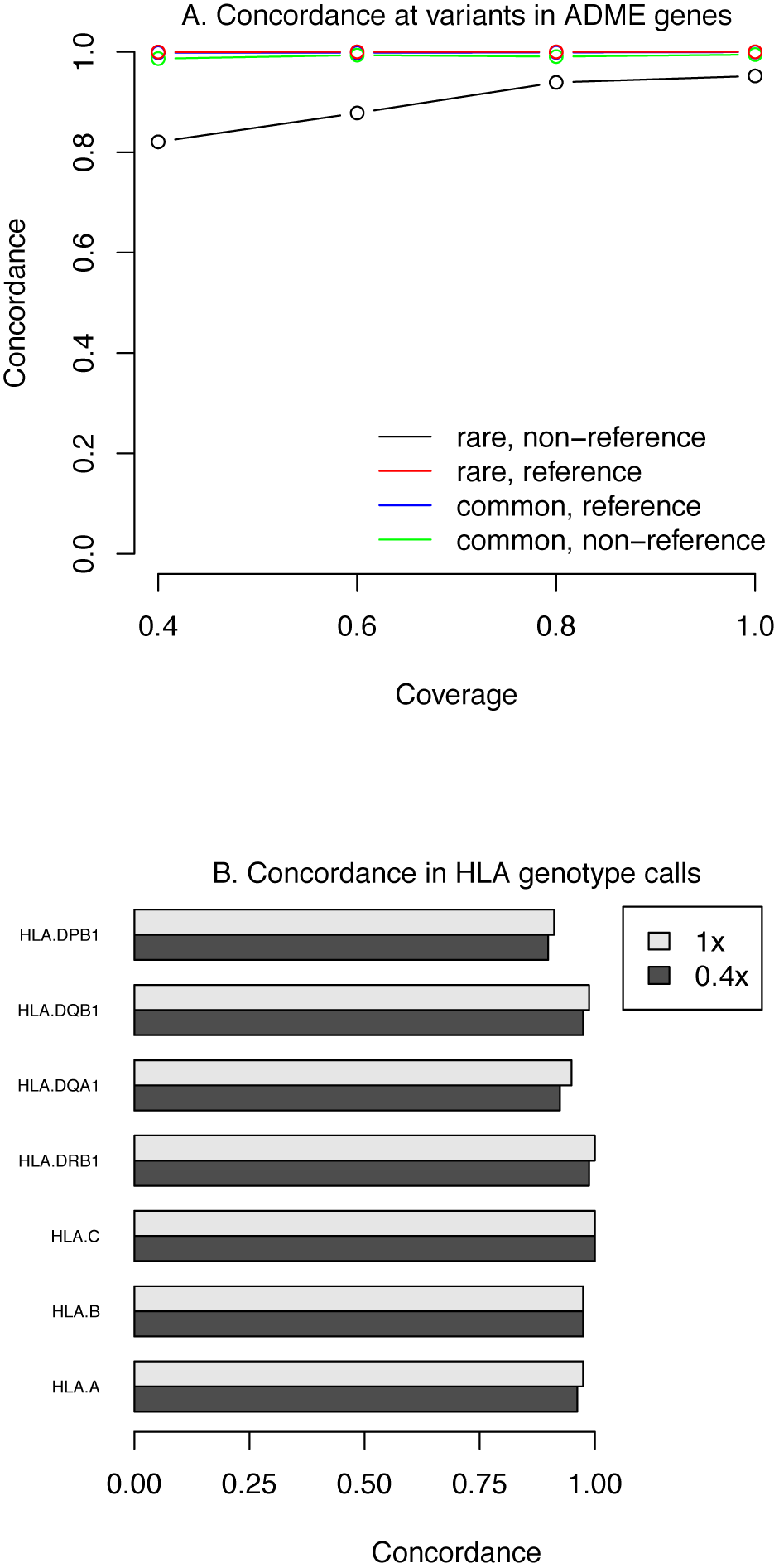
Genotype concordance across platforms at specific variants relevant to pharmacogenomics. **A.** Concordance at SNPs in ADME genes. Variants were classified as “rare” if the minor allele was present in five or fewer copies in the sample (corresponding to an allele frequency of about 3%. Concordance rates are split according to the genotype calls on the PRMA, which was considered “truth”—reference concordance is at variants where the PRMA is homozygous reference, and non-reference concordance is for all other sites. **B. Concordance in HLA genotypes across platforms.** Shown are the concordance rates between sequencing and genotyping array data in imputed HLA genotypes. Concordance is shown for 0.4x and 1x sequencing.

### Genotype concordance at HLA

Apart from ADME genes, another important locus in PGx is the MHC region. We imputed four digit HLA alleles from both the PMRA and sequencing data using HIBAG (Zheng et al., 2014), and assessed the concordance across the two platforms at each of the seven HLA genes assessed by HIBAG. (Figure 1B). There was little variation in imputed genotype concordance across levels of sequencing coverage, and with the exception of the gene DPB1, concordance was above 95%.

For samples where we saw consistent discordance for a given gene between the platforms, we then generated gold standard HLA genotype calls (Methods). A total of 15 HLA genotype calls in 12 samples were retested in this manner. The correct calls were obtained at 7/15 genotypes from the PMRA, and 6, 7, 7, and 8/16 genotypes after imputation from 0.4×, 0.6×, 0.8×, and 1× sequencing, respectively.

### Imputation quality and comparison

Finally, an important metric of how well a technology assays known polymorphisms in the genome is the squared correlation between imputed genotype dosages and the true genotypes (known as “imputation r^2^”). Intuitively, if the researcher has a flat prior on where in the genome to look for an association between a genetic variant and a trait, the average squared correlation is a measure of the power of the study.

We computed this metric for different levels of sequencing coverage by correlating the imputed allelic dosages with directly genotyped sites. We computed this same metric for the genotype data by using the leave-one-out r^2^ at genotyped sites computed by minimac2. At common SNPs, the average r^2^ obtained from the genotyping array was 0.9 (Figure 1), consistent with previous reports from a European population (Nelson et al., 2017). For the sequencing data, this metric varied from 0.9 (for 0.4× coverage) to 0.96 (for 1× coverage).

To investigate the effect of the choice of imputation reference panel on imputation performance on the sequencing data, we performed a head-to-head comparison between using the 1000 Genomes and a subset of the Haplotype Reference Consortium haplotypes (McCarthy et al., 2016) as reference panels using the above methodology (*i.e.*, by treating the array data as “truth” and comparing overlapping sites between the reference panel and the array sites). Using the HRC dataset as the imputation reference panel yielded marginal increases in average r^2^ values in all MAF bins but the lowest, where it suffered a decrease of about 0.036 as compared to the 1000 Genomes imputed sites in the same bin. The exact details and further discussion on this particular comparison can be found in the accompanying Supplementary Materials.

**Figure 2:**
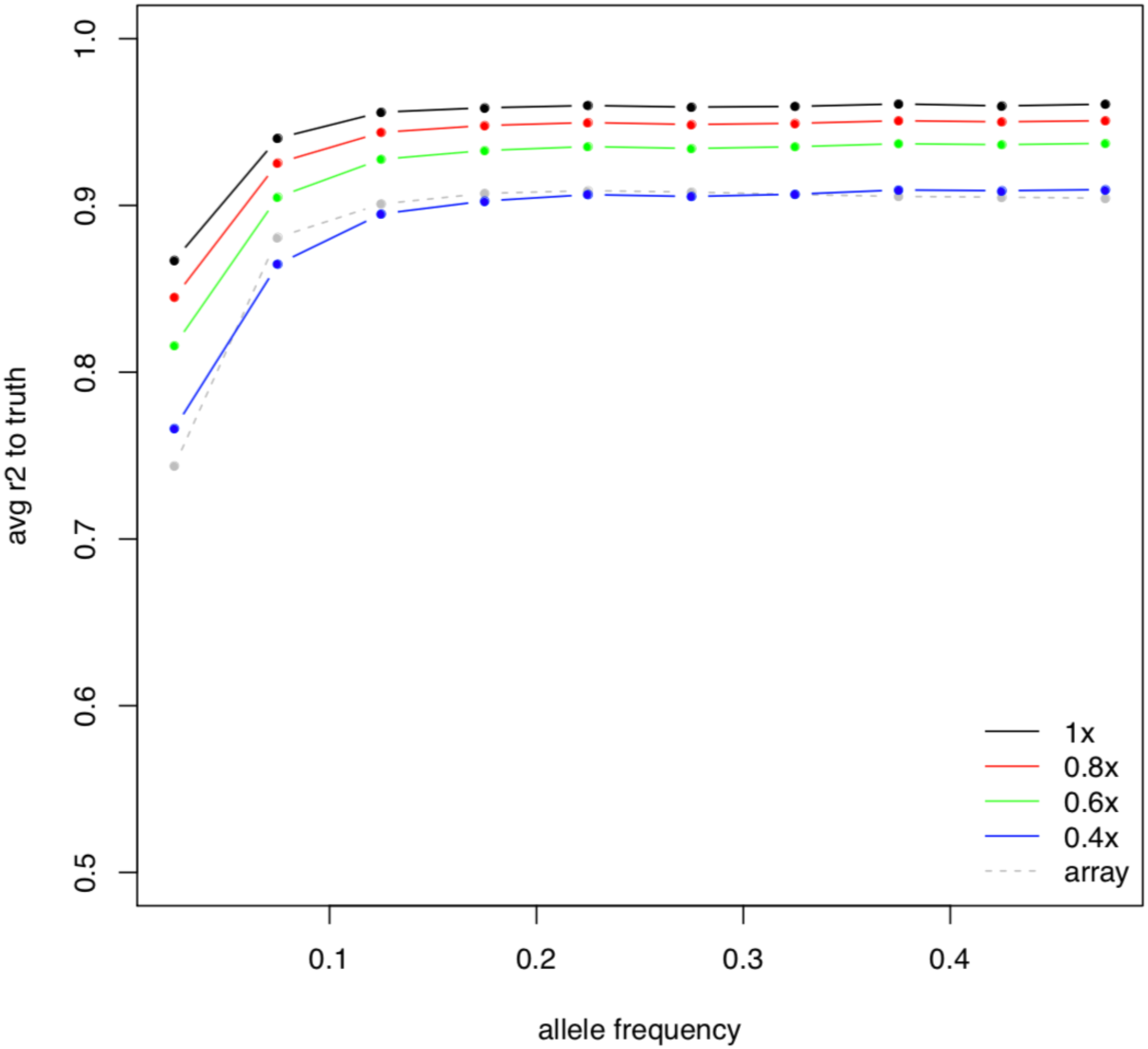
Comparison of imputation quality across platforms. Alleles were binned according to their minor allele frequency (as measured on the genotyping array) and imputation r^2^ averaged across all variants in the bin. For sequencing data, the array data was treated as ‘truth’ and imputation r^2^ computed by correlating imputed dosages to array genotypes. For the array data, imputation r^2^ for all genotyped variants was computed using a leave-one-out procedure implemented in minimac2.

## Discussion

In this paper, we performed a direct comparison between low-pass sequencing (combined with imputation) and a commonly-used genotyping array for the purposes of trait mapping in pharmacogenetics. Overall, genotype calls across the two platforms were highly concordant. For the purposes of trait mapping, low-pass sequencing above a sequencing coverage of 0.4× had higher imputation accuracy than the genotyping array, indicating a corresponding increase in power.

It is worth noting that the cost of sequencing is declining rapidly; if sequencing a human genome to 30× coverage costs $1,000, then the cost of sequencing a human sample to 0.4× coverage is around $13. The key components of cost in a low-pass sequencing assay then become sequencing library preparation and analysis. As the costs of sequencing continue to drop, the importance of these latter costs will continue to grow.

## Methods

### Genotyping

DNA samples were genotyped by BioStorage Technologies/Bioprocessing Solutions Alliance, Brooks Life Sciences (Piscataway, NJ, USA) using the Affymetrix Axiom PMRA.

Prior to genotype imputation, variants in each GWAS dataset were excluded using standard Affymetrix QC thresholds for the PMRA, if there were deviations from Hardy-Weinberg proportions within subgroups of any given ancestry or showed gross and irreconcilable differences in alleles or allele frequency with reference panel genotypes from the HapMap or 1000 Genome projects. Standard Affymetrix array QC sample level thresholds were also applied prior to imputation. (http://www.affymetrix.com/support/downloads/manuals/axiom_genotyping_solution_analysis_guide.pdf)

### Imputation of PMRA data

Genotype imputation for genetic variants that were not directly genotyped (“untyped variants”) was performed using a cosmopolitan haplotype reference panel from the 1000 Genomes Project [The 1000 Genomes Project Consortium, 2015], and using Hidden Markov Model methods as implemented in MaCH and minimac [Li, 2009; Howie, 2012].

### HLA genotyping

High resolution HLA genotyping was performed at BioStorage Technologies/Bioprocessing Solutions Alliance, Brooks Life Sciences (Piscataway, NJ, USA) using the Thermo Fisher AllSet+ Gold SSP High-Resolution HLA kit for HLA-A, HLA-B, HLA-DRB1, HLA-DQB1 and HLA-DPB1 following the manufacturer’s instructions.

### Sequencing

Sequencing libraries were prepared from DNA using the KAPA Library Preparation Kit by Roche and sequenced on an llumina HiSeq 4000 instrument. Sequencing reads for each sample were aligned to the genome using bwa mem (Li and Durbin, 2009), and sequencing reads were randomly sampled to obtain different levels of sequencing coverage. Imputation of genotypes from sequencing data was done using loimpute v. 0.18 by Gencove, Inc. (New York, NY) to a reference panel comprising a subset of the 1000 Genomes Phase 3 (described in more detail in the Supplementary Materials).

### Imputation of sequencing data

Imputation was performed using an implementation of the Li and Stephens model (Li and Stephens, 2003), described in more detail in the Supplementary Note.

