## Supplementary Materials for "Comparing low-pass sequencing and genotyping for trait mapping in pharmacogenetics"

We compared the performance of the imputation of low-pass sequencing data up to two different reference panels with all other parameters held equal, using the performance metrics described in the caption of Figure 2 in the paper. Specifically, we put identically downsampled FASTQ files through Gencove, Inc.'s in-house pipeline and imputed up to (1) the Haplotype Reference Consortium Release 1.1 dataset (accession number EGAD00001002729), and (2) the 1000 Genomes Phase 3 release, trimmed to sites with a minor allele count of >=3. Note that Figure S1 below differs slightly from Figure 2 in the main manuscript due to slight differences in preprocessing and downsampling.

Figures S1 and S2 show the resulting average $r^2$ values in the corresponding allele frequency bins, and Tables S1 and S2 the corresponding values. Given that the HRC dataset is a superset of the 1000 Genomes Phase 3, one would *a priori* expect imputation performance with that dataset to be superior across all allele frequency bins. Somewhat surprisingly, while we did indeed observe a marginal increase in average $r^2$ in every bin but one, imputation on the HRC dataset resulted in lower imputation $r^2$ at the very lowest allele frequency bin – a decrease in the average $r^2$ of about 0.036. We hypothesize that this is an artifact of the strictness in the filtering of the reference panels. Specifically, while the trimmed 1000 Genomes panel contains around 37.5M sites, only around 26.5M of these sites are also present in the HRC (which in total contains around 39M sites despite consisting of an order of magnitude more individuals). We speculate that some of these 11M sites filtered out of the HRC are in linkage disequilibrium with sites genotyped on the array. In a low-pass sequencing experiment at (for example) 0.4x coverage, approximately 4.4M of these sites are expected be covered by a sequencing read and thus contribute information useful for imputation, while in the HRC these variants have been filtered out and thus cannot contribute information that can be used in imputation. If variant filtration in the HRC preferentially affected variants of lower allele frequency (as seems likely), this could lead to a systematic loss of imputation quality at rarer variants when using low-pass sequencing data.

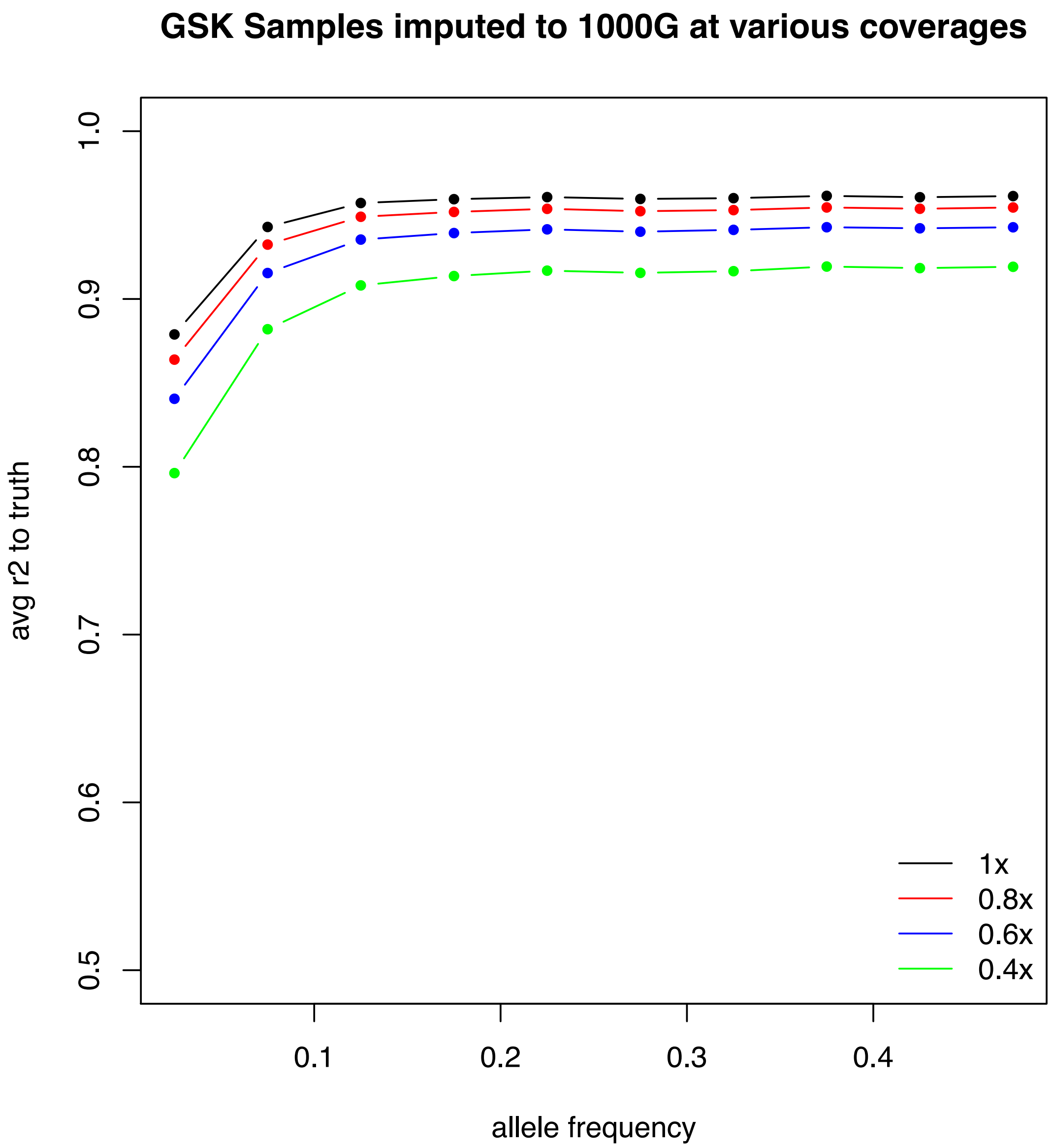


*Figure S1: Comparison of imputation quality at various downsampled coverages using the same metric as in Figure 2 in the main manuscript. The imputation reference panel used was a trimmed version of the 1000 Genomes.*

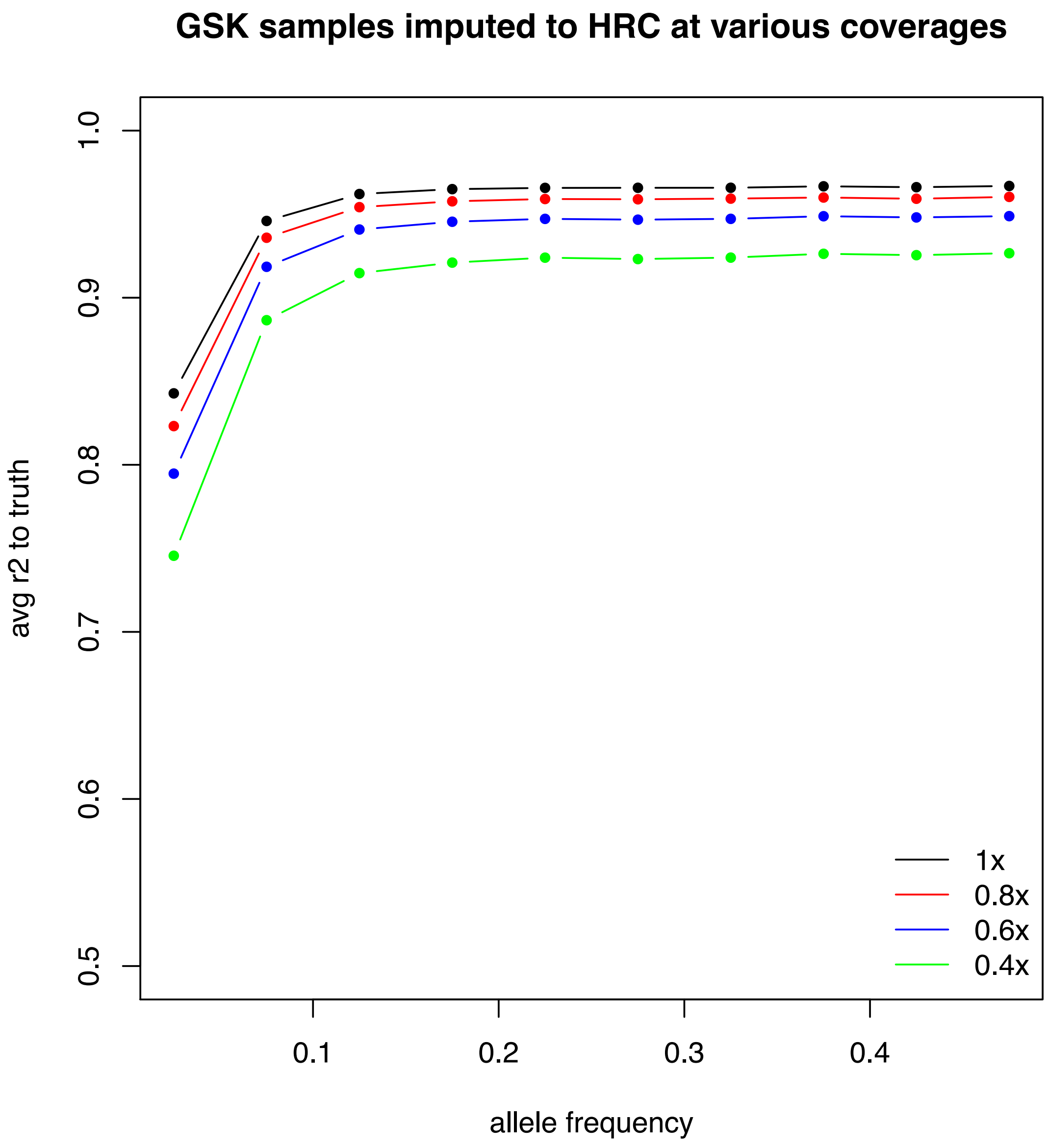


*Figure S2: Comparison of imputation quality at various downsampled coverages using the same metric as in Figure 2 in the main manuscript. The imputation reference panel used was the released subset of the HRC dataset.*

| Allele Frequency Bin | Average $r^2$ to array genotype |
|---|---|
| 0.000-0.050 | 0.842757 |
| 0.050-0.100 | 0.945940 |
| 0.100-0.150 | 0.962060 |
| 0.150-0.200 | 0.964979 |
| 0.200-0.250 | 0.965720 |
| 0.250-0.300 | 0.965779 |
| 0.300-0.350 | 0.965797 |
| 0.350-0.400 | 0.966670 |
| 0.400-0.450 | 0.966143 |
| 0.450-0.500 | 0.966874 |

*Table S1: Values plotted for the data imputed up to the HRC dataset. The data are plotted at the midpoint of each bin. The bins are half-open with the upper end of the interval being closed.*

| Allele Frequency Bin | Average $r^2$ to array genotype |
|---|---|
| 0.000-0.050 | 0.878652 |
| 0.050-0.100 | 0.942976 |
| 0.100-0.150 | 0.957167 |
| 0.150-0.200 | 0.959482 |
| 0.200-0.250 | 0.960772 |
| 0.250-0.300 | 0.959615 |
| 0.300-0.350 | 0.960030 |
| 0.350-0.400 | 0.961296 |
| 0.400-0.450 | 0.960726 |
| 0.450-0.500 | 0.961230 |

*Table S2: Values plotted for the data imputed up to the trimmed 1000 Genomes Phase 3 dataset. The data are plotted at the midpoint of each bin. The bins are half-open with the upper end of the interval being closed.*
