## Supplementary Note for "Comparing low-pass sequencing and genotyping for trait mapping in pharmacogenetics"

### Supplementary Note for: Comparing low-pass sequencing and genotyping for trait mapping in pharmacogenomics

April 12, 2019

This note describes the imputation algorithm for low-pass sequencing data implemented in the software loimpute v0.18 by Gencove, Inc. The basic setup is the copying model proposed by Li and Stephens [2003]. Assume we have a set of $N$ bi-allelic variants that have been genotyped on a set of $M$ phased haplotypes, such that the allele at variant $i$ on haplotype $j$ is $h_{ij}$, coded as a 0 if the allele matches the reference genome and as a 1 if the allele matches the alternate allele at the site. At each of the $N$ sites we additionally have a set of $L$ sequencing reads from the individual whose genotypes we would like to impute, such that the allele at variant $i$ at sequencing read $k$ is $r_{ik}$. We now wish to impute the genotype of the (diploid) target individual at each of the variants.

Now let $X_{i,1}$ and $X_{i,2}$ be the identity of the haplotypes being copied at variant $i$ by the target individual. Similarly to Li and Stephens [2003], the transition probabilities from $X_{i,1}$ to $X_{i+1,1}$ can be modeled as:

$$P(X_{i+1,1} = x' | X_{i,1} = x) = \begin{cases} \exp(-\rho d_j/M) + (1 - \exp(-\rho d_j/M))(1/M) & x' = x \\ (1 - \exp(-\rho d_j/M))(1/M) & \text{otherwise,} \end{cases}$$

where $\rho$ is the population-scaled recombination rate per base (set for all applications here to 0.0001) and $d_j$ is the physical distance between the variants at positions $i$ and $i + 1$. The setup for the second haplotype of the individual is identical.

Now let $Y_i = X_{i,1} + X_{i,2}$. The emission probabilities of each read given $Y_i$ can be modeled as:

$$P(r_{ik} = r | Y_i = y) = \begin{cases} \epsilon & r = 0, y = 2 \\ 1 - \epsilon & r = 0, y = 0 \\ 0.5 & y = 1 \\ 1 - \epsilon & r = 1, y = 2 \\ \epsilon & r = 1, y = 0, \end{cases}$$

where $\epsilon$ is the sequencing error rate (set for all applications here to 0.001).

The posterior probability of each genotype in the target individual can be estimated using standard methods for hidden Markov models.

**Computational complexity.** Note that this algorithm has time complexity of $O(NM^2)$, which makes computation infeasible even for reference panels of modest size. To mitigate this, we need to choose a small set of plausible haplotypes to run the full algorithm. To do this, we identify a set of haplotypes that share rare variants with the target individual, and then use only those haplotypes in a given region. Specifically, in some region, consider all of the genetic variants with a frequency in the reference panel less than some threshold $f$ (for our purposes, we set $f$ to 0.05). Now for each haplotype, compute the statistic $f_M/f_T$, where $f_M$ is the number of alternative alleles at these sites where the haplotype and at least one sequencing read from the target individual match, and $f_T$ is the count of the sites where the haplotype carries the alternative allele at the site. We rank haplotypes according to this statistic, and choose the top $M'$ haplotypes to run the full model. For all applications, we set $M'$ to 100.
